# Taking extreme measures: A quantitative study of multiple stress interactions at the limits of life

**DOI:** 10.1101/2020.01.13.905562

**Authors:** Rosie Cane, Charles Cockell

## Abstract

Environments exposed to simultaneously occurring extremes are prevalent in the natural world, yet analysis of such settings tends to focus on the effect of single environmental stresses. In this study, quantitative multiplicative and minimising models previously used to study nutrient limitation were applied to the growth of the hydrothermal vent-dwelling organism *Halomonas hydrothermalis* when subjected to combined nutrient limitation and NaCl-salt stress. Results showed an interactive effect from both salt and nutrient stresses under optimal conditions. However, the fit became more non-interactive as salinity is increased; at which point NaCl-salt had a more dominating effect on growth than inorganic phosphate (P_i_). We discuss biochemical hypotheses to explain these data. This work shows that models developed to understand nutrient limitation can be used to quantify and separate the contributions of stresses under other physical and chemical extremes, such as extreme salinity, and facilitate the development of biochemical hypotheses of how extremes may be influencing cell physiology.

**Importance:** Very few environments in the natural world are exposed to just one extreme or stress at a time. To understand life’s ability to survive in multiple-extreme environments, we must be able to quantify how different extremes interact. Using methods developed for the study of multiple nutrient limitation, this study uses kinetic growth models to investigate at the effect of extreme environments on bacterial growth. Results show that closer to the extremes of life, individual stresses dominate growth; whereas under optimal conditions there is a multiplicative effect from both salt and nutrient stresses. This approach offers a new way to quantify and potentially understand and develop hypotheses for how life operates under multiple extremes.

## Introduction

Micro-organisms are able to tolerate a variety of stresses which has enabled them to colonise many environments once thought to be uninhabitable, known as extreme environments [1]. For decades, laboratory studies into extreme environments have tended focused on single extremes, such as temperature, pressure, pH and salinity [2]. Although this provides an insight and enhanced understanding of the absolute limits to life under a given physical or chemical extreme [3], very few environments in the natural world expose life to just the one stress [2]. Most microorganisms are exposed to multiple extremes and yet there have been relatively few studies conducted to research them. Those studies that do investigate multiple extremes tend to measure microbial growth parameters, but provide little analysis of how the component extremes influence the combined extreme responses.

However, methods have been used in other areas of microbiology to quantify the relative contributions of individual stresses to a combined stress response. This is particularly the case in the study of nutrient limitation. The concept of co-limitation has been explored by several studies in the fields of oceanography and food industry [4],[5], reviewing the effects of nutrient co-limitation on bacterial growth. Furthermore, studies such as Darvehei et al., 2018 [6] explore models developed on the behaviour of archaea cultures, a domain of life often encountered in multiple extreme environments, built to account for individual effects of light, nutrients, temperature, pH and dissolved oxygen; highlighting the significant need of a method to explore multiple-stresses.

Synchronously occurring stresses were first explored over 40 years ago [7], introducing the terms ‘interactive’ and ‘non-interactive’ to describe the relationship between two nutrient stressors. An interactive response illustrates both substrates to affect the overall growth rate of an organism in a synergistic manner, and a non-interactive response shows that an organism can only be limited by one substrate at a time.

Previous attempts to quantify interactions between stresses generally focus on models specific to particular stressors. For example, Ross et al 2003 [8] employed a square root model with stress-specific parameters to investigate the effect of temperature, water activity, pH and lactic acid concentration, while Leiss et al 2015 [9] used a ‘Stress Addition Model’ model to study stressors in aquatic systems. These are limited in use since they do not allow for the examination of a range of different stresses with the same approach. Advancing on this work, the aim of this study is to produce a simple quantitative method that can be applicable to all stresses, and not limited by specific functions.

In this study, we investigated how nutrient limitation concepts can be translated into the study of the effects of multiple extremes on microorganisms. We use the Monod Multiplicative Model [10] and Liebig’s Law of the Minimum [11] as a basis – both simple and continuous algebraic relationships which lend themselves readily to theoretical analysis [7]. Monod kinetics provide an interactive model, allowing for the investigation of the multiplicative effects of stresses, and Liebig kinetics provide a non-interactive model, allowing for the investigation of a ‘minimising’ approach, whereby one stressor dominates the stress response. These two kinetic growth models have been widely compared in a nutrient-limitation context ([12]-[14] etc). Here we use them to quantify effects of combined extremes.

## Results

### Growth results of *Halomonas hydrothermalis* under Salt and P_i_ stresses

Results investigating the growth of *Halomonas hydrothermalis* under salt and P_i_ stresses show how growth rate varies under differing concentrations of each stressor. **Figure 1** shows the effect of low, optimum and high salinities (defined here as 0, 3-4 and 8% [wt/vol] respectively), each at low (0.3 M) and optimum (varying with NaCl concentration) P_i_ concentrations on growth rate. In this study, optical density data was transferred to Microsoft Excel where growth curves were produced, using the exponential phase to produce growth rates, acquired using equation 1 (See Figure 2 (b)).

**Figure 1.**
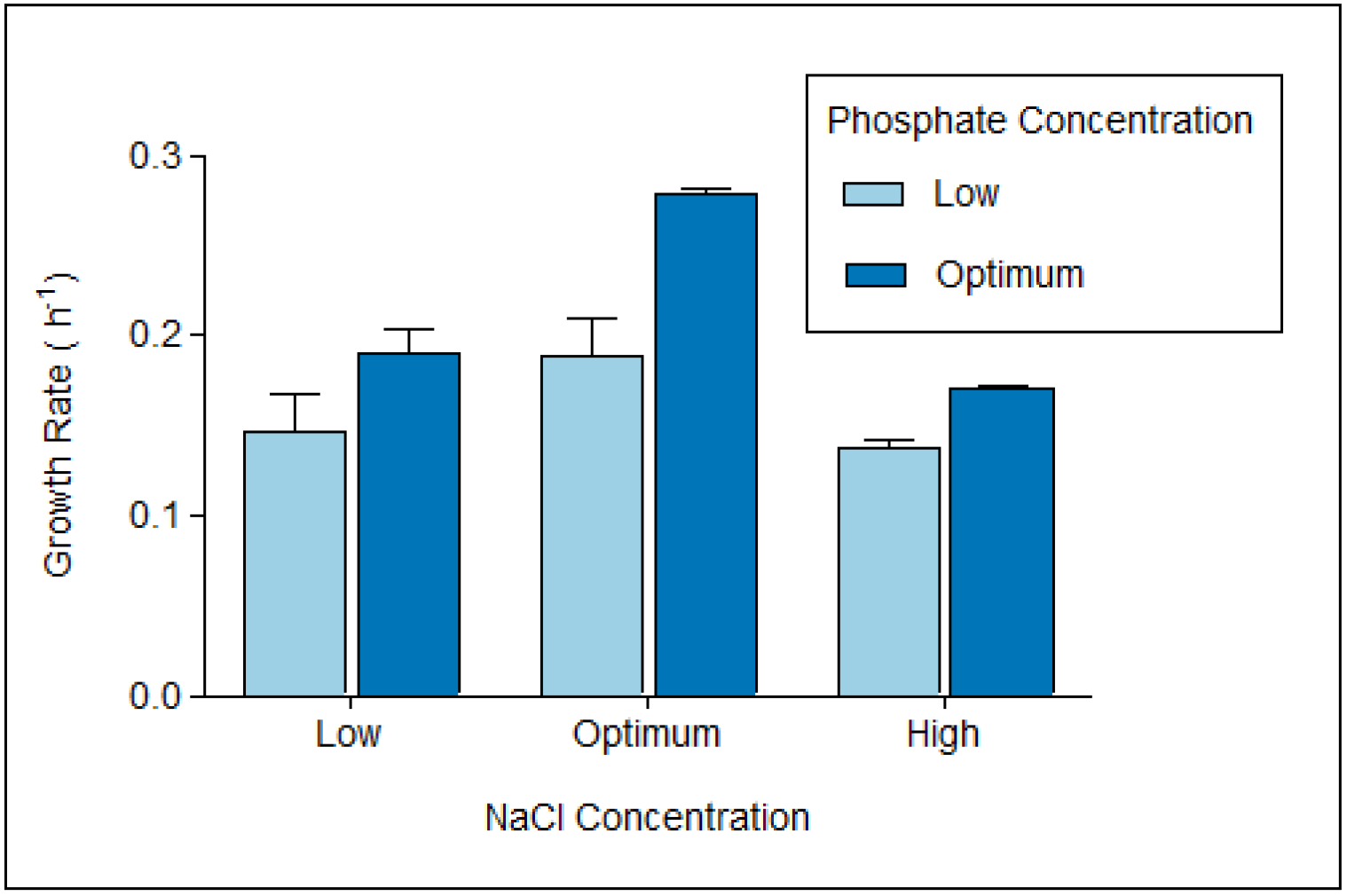
Growth rates of the organism *Halomonas hydrothermalis* under combined NaCl and P_i_ limitation. Growth rates were calculated from growth curves as described in Materials and Methods. Data are presented as untransformed means ± standard errors of the means (SE).

**Figure 2.**
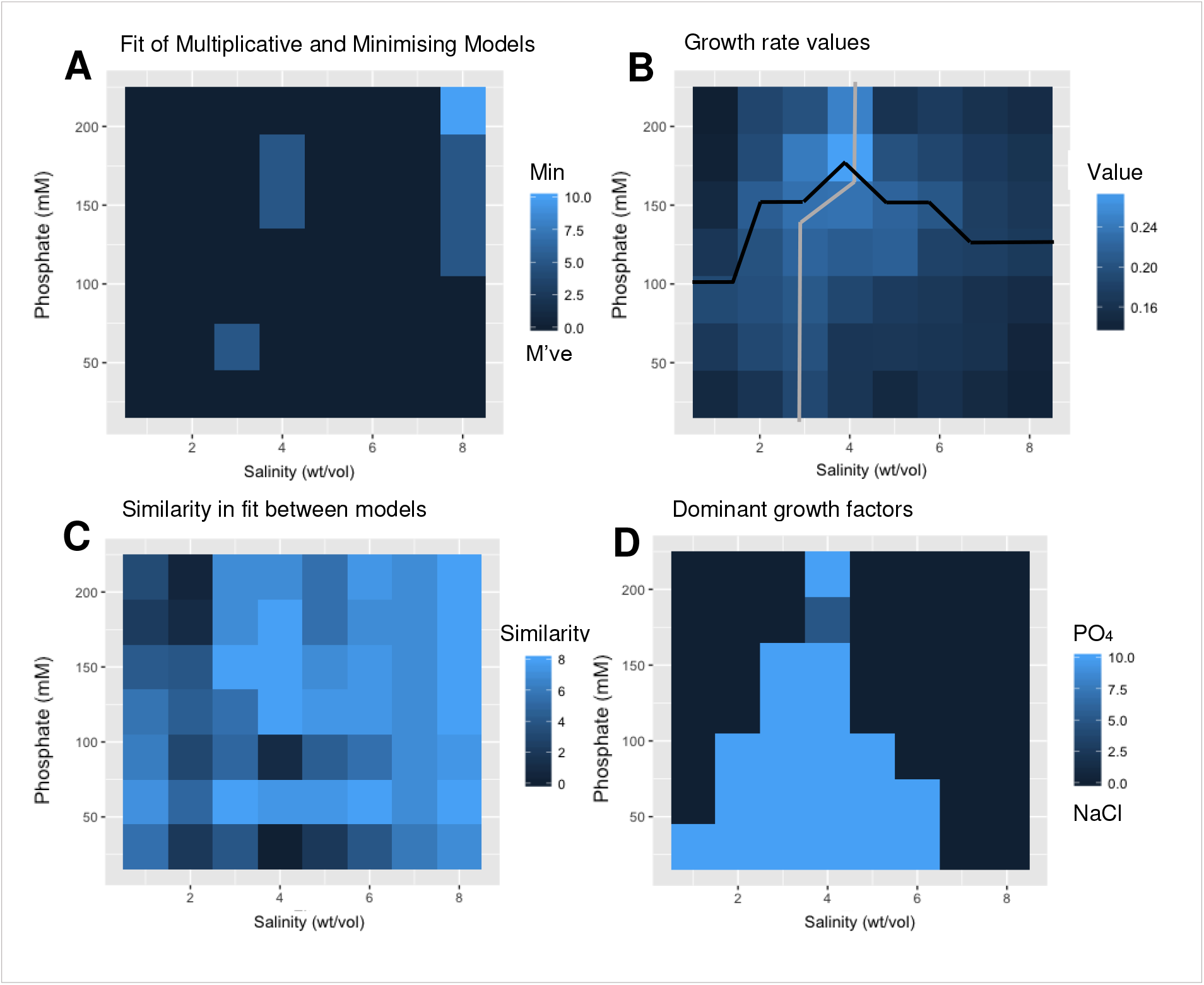
Fitting minimizing and multiplicative models to life in extremes. Heat maps showing quantitative analysis of growth under combined salt and P_i_ stresses. (A) Fit of multiplicative to minimising model; best fit to multiplicative model shown in dark blue and minimising light blue. (B) Growth rate (μ) under different combinations of salt and P_i_, showing optimal salt concentrations (grey line) and optimal P_i_ concentrations (black line). (C) Similarity between fit of multiplicative and minimising models – light blue = similar fit and dark blue = large difference in fit. (D) Whether salt or P_i_ dominated growth - light blue = P_i_, dark blue = salinity.

Growth rate results show a general positive correlation between growth rate and increase in NaCl up until an optimal growth rate was reached, followed by a negative correlation as growth rates decreased. Optimum growth was achieved at 4% (wt/vol) NaCl and 0.18 M P_i_. Under optimal conditions for both stressors, the organism exhibited an average growth rate of 0.278. No growth was observed below 0.3M P_i_ or above 8% (wt/vol) NaCl at non-optimal P_i_ concentrations.

A multiple-regression ANOVA was implemented (**Table 1**), with a subsequent box-cox transformation to reduce data skew. After accounting for variation in time, statistical tests showed that all factors were significant. Following this, a Post-hoc Tukey Honest Significance Difference (HSD) test was carried out to identify the interaction between stresses, which produced Least Squares Mean values, indicating pair-wise similarities between low and high salt concentrations. Homogeneity of variables was also assessed with an *F*-test, giving a *p*-value of <0.05. Following this, unequal variances were taken into account for the *t*-test.

**Table 1.**
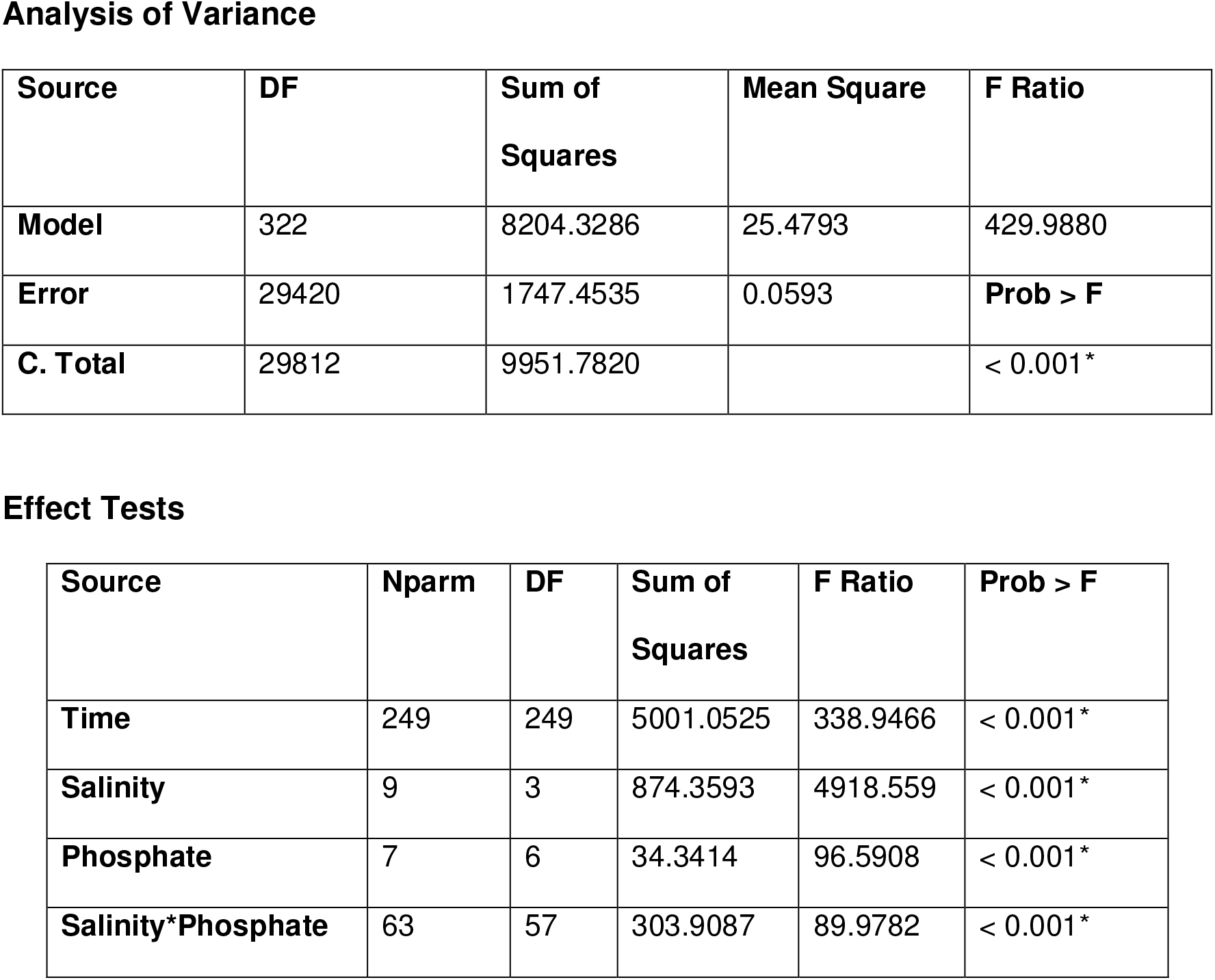
Results of ANOVA investigating effects of phosphate and salinity over time on growth of the organism *H. hydrothermalis*.

Analysis of Variance
| Source | DF | Sum of Squares | Mean Square | F Ratio |
| --- | --- | --- | --- | --- |
| **Model** | 322 | 8204.3286 | 25.4793 | 429.9880 |
| **Error** | 29420 | 1747.4535 | 0.0593 | **Prob > F** |
| **C. Total** | 29812 | 9951.7820 |  | < 0.001* |

| Source | Nparm | DF | Sum of Squares | F Ratio | Prob > F |
| --- | --- | --- | --- | --- | --- |
| **Time** | 249 | 249 | 5001.0525 | 338.9466 | < 0.001* |
| **Salinity** | 9 | 3 | 874.3593 | 4918.559 | < 0.001* |
| **Phosphate** | 7 | 6 | 34.3414 | 96.5908 | < 0.001* |
| **Salinity\*Phosphate** | 63 | 57 | 303.9087 | 89.9782 | < 0.001* |

### Fit of quantitative methods

To present the data obtained in this study, we have organised it into a four-plot schematic. The fit of the two quantitative methods is visualised as four heat maps (Figure 2); a) the fit of the multiplicative and minimizing models, b) growth rate values, c) the similarity of fit between these two models and d) whether salt or P_i_ dominates growth.

Map (a) shows fit of the multiplicative (dark blue) and minimizing models (light blue). The majority of the combinations of the two stresses have a better fit to the multiplicative model with the exception of: 3% (wt/vol) NaCl and 0.06 M P_i_, 4% (wt/vol) NaCl and 0.25-0.18 M P_i_ and at a combination of high salt and ≥ optimal P_i_ concentrations. Map (b) presents growth rate values at different concentrations of P_i_ and salinity. It can be seen that the highest growth rates (light blue) occur under optimal salt and P_i_ concentrations, and that growth rate decreases (darker blue) as high and low salt stresses are reached. Optimal P_i_ concentrations (grey line of best fit) varied with salinity, starting at 0.09 M at 1%, increasing to 0.18 M at optimal salt, and then decreasing to 0.12 M at 8% (wt/vol). Optimal salinity (black line of best fit) was shown to vary from 3 to 4% (wt/vol), with highest growth observed under optimal P_i_ concentrations.

Similarity between the fit of both multiplicative and minimizing models can be seen in Map (c). A highly similar fit, i.e. both multiplicative and minimising models closely fit (light blue) is generally observed at the edges of the heat map at more extreme salinities, and more centrally under optimal salt and P_i_ concentrations. Map (d) displays whether P_i_ or salinity is having the greatest effect on growth at any given combination of the two stresses. Given that the majority of fit is to the multiplicative model, this shows where one stress is having more of an effect than the other. It can be seen that at high P_i_ and both high and low salt concentrations that the effect of high/low salinity has a greater effect on growth. At lower P_i_ concentrations and optimal salinities, P_i_ generally affects growth more than salinity.

At low salt concentrations (1-2% [wt/vol]), growth is mainly dominated by the effects of a lack of NaCl, with fit corresponding to the multiplicative model. However, the similarity between fits of the two models at 1% (wt/vol) salinity and 0.03-0.12 M P_i_ is very high. Additionally, at optimal salt concentrations (3-4% [wt/vol]), P_i_ affects growth the most at lower P_i_ concentrations, and salt affects growth the most at higher Pi concentrations. Figure 2 (A) shows nearly a complete multiplicative fit, aside from at 4% (wt/vol) and 0.15-0.18 M P_i_, and also 3% NaCl and 0.06 M P_i_, where the fit is very similar between multiplicative and minimizing. At high salt concentrations (5-8% [wt/vol]), as salinity increases, fit becomes closer to that of the minimizing model. When 8% (wt/vol) NaCl is reached, a similarity in fit between both multiplicative and minimizing models can be observed between 0.12, 0.15 and 0.18 M of P_i_. A fully-minimizing fit is observed at 8% (wt/vol) NaCl and 0.21 M of P_i_.

## Discussion

A large number of multiple-extreme environments exist in the natural world, yet very little has been done to investigate interactions between extremes. In particular, studies tend to focus on the physiological effects of one extreme or combined extremes without attempting to understand quantitatively how those individual extremes interact to regulate growth. In this study, we carried out experiments to investigate whether models developed to understand the interactions of nutrient stress could be applied to quantifying the contributions of stresses that include the physical and chemical extremes of life. Here we studied the effects of salt (NaCl) stress and P_i_ limitation on bacterial growth. Both NaCl and P_i_ can affect growth individually, but when combined an interaction can be anticipated between the two stressors [15].

### Experimental growth rates

Lowest rates of growth can be observed under the highest and lowest salinities at non-optimal P_i_ concentrations. This occurs when these extremes are approached, more severe limitation occurs and there is less growth. However, at optimal P_i_ and salt concentrations, where there are minimal stresses present to hinder the growth of bacterial cells, rates of growth are much higher.

### Multiplicative and Minimizing growth dynamics

The majority of combinations of P_i_ and salt stresses have best fit to the multiplicative model. We interpret this to show that most combinations of salt and P_i_ limitation stress are interactive when not at the absolute limit. This can be understood at the biochemical level. We would expect that when an organism is not growing at the limit of the imposed stresses no stress would dominate and determine the threshold of growth. Rather, away from the extreme edges any imposed stress will add to the total energetic demand and/or biochemical response in a multiplicative way.

At 4% (wt/vol) and 0.15-0.18 M P_i_ there is a high similarity between fit to multiplicative and minimizing models. These conditions represent optimal salt and P_i_ concentrations, and as no stresses are acting upon the bacterium in this instance, an exact fit to either model should not be anticipated.

### NaCl-salt and P_i_ stress effect on growth at extremes

At low salt concentrations, growth is observed to be mainly dominated by salinity. This fit corresponds with the multiplicative model, and although salt is observed to have a more prevalent effect than P_i_, both stresses are still affecting growth.

When at optimal salt concentrations, although still under multiplicative kinetics, P_i_ has a greater effect on growth than salt. In this instance, salt would not be anticipated as a limiting factor as the bacterium maintains optimal salt concentrations for cell regulation.

At high salinities, the effect of high salt concentrations dictates growth. As salinity increases, salt has a greater effect on bacterial growth, until the fit is best to the minimizing model at 8% (wt/vol) and 0.21 M P_i_. *Halomonas* and other members of the *Halomonadaceae* family have been shown by Gunde-Cimerman et al, 2018 [16] to use organic solutes for osmotic stabilization and to maintain low cytoplasmic ionic concentrations under highly saline conditions – a system called the ‘compatible solute strategy’ [17]. If this is the case for *Halomonas hydrothermalis*, an explanation of the minimizing fit at high salt and P_i_ conditions could be energetic constraints. Under these conditions, energy demands might be dominated by osmotic regulation to the point where this demand dominates cell responses. In support of this, Rai et al, 2005 [18] found that P_i_ uptake in the cyanobacterium *Anabaena doliolum* was significantly reduced with an increase in salinity, suggesting that P_i_ could not be efficiently transported into cells. The study concluded that energy constraints that may have caused an absence of strategies to uptake P_i_ under high salt concentrations, as P_i_ uptake is an energy-dependant process. Thus although under these conditions, phosphate would be limiting, high salinity would ultimately be the controlling factor in growth.

Another explanation could be nutrient over-exposure under high salt concentrations. Previous studies have focussed on P_i_ limitation at high salinities, but not on high concentrations of the nutrient. The general bacterial response mechanism for P_i_ uptake in conditions with an abundance of P_i_ is the P_i_-Inorganic Transport (Pit) model [19]. An excess in P_i_ in highly saline environments can have an adverse effect on regulation of P_i_ metabolisms via the Pit model [20], and indeed very few bacterial strains with P_i_ solubilizing abilities can function well at high concentrations of NaCl [21]. High salt concentrations are effectively cutting off the ability of the organism to regulate P_i_; leading to salinity dominating growth through the minimizing model.

Yet another hypothesis could be linked to the uptake of potassium (K). In saline environments, some cells accumulate K^+^ ions to balance osmotic concentrations inside and out of cells – the ‘salt-in strategy’ [22]. In addition, P_i_ transport systems are dependant upon the presence of K^+^ [19]. This indicates that at higher salt concentrations, cells may be taking up K^+^ for osmoregulation processes, and depleting available P_i_. Uptake of potassium ions might change the ionic associations of K^+^ with P_i_ ions, indirectly affecting the ease by which P_i_ can be taken up by cells. If the presence of high Na^+^ and Cl^−^ concentrations influences P_i_ concentrations, then at extremes of NaCl, it might be observed to dominate growth.

These data show how multiplicative and minimising models can be used to generate biochemical hypotheses, which could be tested with further laboratory experiments. In this case, more needs to be known about how P_i_ and NaCl interact and the molecular mechanism(s) of P_i_ solubilization in salt-stressed environments to explain the kinetic data obtained.

## Conclusion

Although a great many studies investigate the effects on one extreme on microbial growth or the combined effects of multiple extremes on growth, here we have used kinetic models to show how concepts borrowed from studies on the effects of nutrient limitation can be expanded to study the interactions of multiple extremes and to tease their effects apart. We have shown that by not only measuring growth rates, but by defining the role of multiplicative or minimising interactions at extremes, not only can the interactions of combined extremes be quantified, but that they lead to the development of new hypotheses. In this case, the hypotheses offered to explain the growth minimising domination of salt stress over P_i_ stress at high salt and P_i_ levels could be investigated by proteomics, for example. We conclude that the methods developed to examine nutrient limitation stresses offer a powerful way to quantify and examine the behaviour of life in extreme environments.

As well as informing an understanding of how life adapts to extremes, this work has implications for the assessing the habitability of extra-terrestrial environments. The models described here might be used to investigate the growth of organisms grown under extremes relevant to extraterrestrial environment to develop hypotheses on how much extremes would be expected to limit life, for example contaminant microorganisms transferred to other planetary bodies.

Multiple-extremes have been quantified a new method of looking at extremes which allows us to develop hypotheses for biochemical effects of multiple extremes.

## Materials and Methods

### Strain and growth conditions

The bacterium *Halomonas hydrothermalis* was selected for this study. It was isolated from an extreme hydrothermal environment [23] and is adapted to a wide range of stresses [15]. From the family *Halomonadacacae*, popular in research on osmotic adaptation due to their moderately halophilic nature [16], this bacterium has well-documented boundaries of growth, exhibiting cell division across temperatures of 2 to 40°C (optimal growth reported at 30°C), total salt concentrations of 0.5% to 22% (wt/vol) (optimal range of 4% to 7% (wt/vol)), and pH values of 5 to 12 (optimal range of 7 to 8) [15],[23]. Although the salt-tolerance of this organism has been constrained, nothing is known about required nutrient levels for cell division to occur.

The organism was cultured in a marine-minimal media as described by Ostling et al., 1991 [24], with the following recipe: (a) 920 ml 1.1 × NSS (artificial seawater) [25]; (b) 10 ml 132 mM K_2_HPO_4_; (c) 10 ml 952 mM NH_4_CI (pH 7.8), and the following sterile filtered solutions; 10 ml 0.4 M tricine, 1 mM FeSO_4_ • 7 H_2_0 (pH 7.8); (e) 40 ml 1 M MOPS (potassium morpholinopropane suifonate, pH adjusted to 8.2 with NaOH); (f) 10 ml of 20% glucose stock solution.

### Choice of stress conditions

Laboratory experiments were conducted to study the effect of NaCl-salt and P_i_ stressors and co-stressors on the growth of the organism *Halomonas hydrothermalis*. These measurements allowed us to test at the physiological level whether there is an interaction between nutrient limitation and salinity, and how this was affected as extremes of life were approached [26]. Salt limitation was chosen because of the prevalence of salinity as a variable factor in many extreme environments, including evaporites, volcanic pools, and hydrothermal vents [27],[28] and is of considerable interest in astrobiology and the assessing the habitability of extraterrestrial environments [29], since saline environments have been observed elsewhere in our Solar System [30]. In addition to being confronted with physical and chemical extremes, microbes are often limited for nutrients. Phosphorus, in the form of inorganic Pi (P_i_), is limiting in many natural ecosystems and is one of the most important macronutrients for maintaining cellular functions [31]. Both factors have prevalence in the habitability of Earth’s oceans, and could be extended to understanding the habitability of aqueous environments beyond the Earth. Furthermore, the interaction of these two extremes is virtually unknown outside of soil ecology, and following this, this study aims to provide an insight into synergy between salinity and P_i_ concentration in other extreme environments, such as hydrothermal environments from which our model organism was isolated.

### *Halomonas hydrothermalis* growth curve experiments

An exponential-phase culture was obtained by growing the bacterium in marine-minimal media and incubating at 30°C. Aliquots were prepared (50% [vol/vol] glycerol) for storage at −80°C. The stored cultures were used to inoculate marine-minimal agar (3.5% [wt/vol] NaCl), which was incubated at 30°C for 48 h and stored at 4°C until use. Cultures were left overnight in a shaking incubator at 30° C and prior to experiments, cells were washed to remove any residual P_i_. This was done using a Fisher Scientific TopMix FB15014 vortex to mix the culture, then washing cells with P_i_ -free media using a Thermo Electron Corporation Inc MicroCL17 centrifuge for 2 minutes at 5-G to leave a pellet. The supernatant was removed and replaced with phosphate-free marine minimal media. This process was repeated 3 times. Cells were then diluted to give a cell density equivalent to an optical density at 600nm (OD_600_) of 0.4.

A Greiner-Bio Flat Bottom 96-well plate was used for experimental layout, with varying salinity (0-8% [wt/vol]) and P_i_ (0-0.21 M) across the plate. All experiments were conducted at 30°C. Optical density measurements were carried out over 24 hours using a BMG Labtech SpectroSTAR Nano Microplate Reader (manufacturer and location) taking reads every 1.5 minutes. Growth rates (μ) were calculated using the exponential growth curve phase and measured using Equation 1 [32].

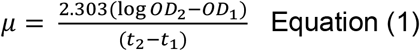

### Statistical Analysis

Interaction between salinity and P_i_ concentrations was investigated using multiple regression analysis. An initial analysis of variance (ANOVA) test was carried out using JMP software (JMP Version 12, SAS Institute Inc, Cary, NC), leading to an effects test to investigate the synergy between the two stresses. Data was Box-Cox transformed to reduce skew in distribution and a Posthoc Tukey Honest Significance Difference (HSD) test was undertaken to analyse the difference in least square means values.

### Data Fitting

Experimental data was split into three data sets: dataset 1, in which growth was influenced by both P_i_ limitation and salinity extremes, dataset 2, where growth is only affected by salinity and dataset 3 where growth is only affected by P_i_ limitation. To test whether the multiplicative or minimizing method exhibited the greatest effect on growth, an unpaired, two sample *t*-test was carried out. This compares a mean and predicted standard deviation for datasets 2 and 3 combined (derived either to fit the multiplicative or minimizing method) with the mean and standard deviation for dataset 1.

To predict a standard deviation for the multiplicative approach, a Gaussian, normal distribution was assumed for simplicity. The product of variables in relation to dataset 2 and dataset 3 was calculated to give a predicted multiplicative variance of the two combined. For the minimizing approach, the distribution of the minimum of random Gaussian variables, moments Y=min(X_1_, X_2_) [33] were applied using probability density function and cumulative distribution functions to predict an overall variance.

For investigations into which of the two stresses had a greater effect on growth, a one-sided students *t*-test was carried out between the two distributions in R-studio. This showed whether the average of the salinity factor for replicates is greater than average of the nutrient limitation factor. Results were visualised using heat maps produced in R Studio; showing the fit of the two methods, growth rate values, which stress dominated at which combination of stresses, and similarity in fit of the two models. The latter was calculated using the difference between *p*-values for the multiplicative and minimizing models, then dividing results into 40 categories between 0 and the maximum difference, 0.0881. Each of these categories was assigned a number between 1 and 10 in increments of 0.25 to visualise the goodness of fit of the two models.

## Acknowledgements

A huge thank you to Dr Rosalind Allen and Dr Rebecca Prescott at The University of Edinburgh for their statistical guidance through this work. Also thanks to the Principal’s Career Development Scholarship at The University of Edinburgh for funding this work. Charles Cockell was supported by Science and Technologies Facilities Council Grant number: ST/R000875/1.

